# Structural Connectivity Selectively Constrains Intrinsic BOLD Timescales through Graph-Smooth Neural Activity

**DOI:** 10.64898/2026.06.14.732146

**Authors:** Ali Bashirgonbadi, Mohamad Reza Salehi, Hamid Soltanian-Zadeh

**Author notes:** Corresponding author: Hamid Soltanian-Zadeh.

## Abstract

Structural connectivity defines the network architecture supporting large-scale brain dynamics, yet how this network constrains the temporal statistics of signals defined on it remains poorly understood. Prior work has reported associations between intrinsic timescales of resting-state fMRI and structural connectivity strength, but it is unclear which signal components primarily drive this relationship. Here, we adopt a graph signal processing framework to analyze intrinsic temporal properties of networked brain signals. Regional Blood Oxygenation Level Dependent (BOLD) activity is modeled as a graph signal supported on the structural connectome and decomposed via graph spectral filtering into low-frequency (structure-coupled) and high-frequency (structure-decoupled) components. Using diffusion MRI–derived structural connectivity and resting-state fMRI from 100 unrelated participants of the Human Connectome Project, intrinsic timescales are quantified using relatively low-frequency power and related to node-wise structural connectivity strength while controlling for regional volume. We show that intrinsic timescales derived from structure-coupled signals exhibit robust positive associations with structural connectivity strength at both group and inter-individual levels, whereas structure-decoupled signals display substantially weaker coupling. Notably, slow structure-decoupled dynamics are preferentially expressed in higher-order association cortex. Graph-spectral null models further demonstrate that these effects critically depend on the empirical organization of the structural network. Together, these results establish a graph-spectral interpretation of structure–timescale coupling, showing that network topology selectively constrains the temporal statistics of graph-smooth neural activity.

**Author Summary:** A fundamental question in network neuroscience is how the brain’s structural connectivity shapes the temporal dynamics of functional activity. Previous studies have shown that brain regions with stronger anatomical connectivity tend to exhibit slower intrinsic activity fluctuations, but the functional signal components responsible for this relationship remain unclear. Here, we combine graph signal processing with analyses of intrinsic BOLD timescales to separate resting-state activity into structure-coupled and structure-decoupled components. Using multimodal neuroimaging data from the Human Connectome Project, we show that the association between structural connectivity strength and intrinsic timescales is primarily driven by structure-coupled, graph-smooth activity. In contrast, structure-decoupled dynamics exhibit substantially weaker dependence on anatomical connectivity, although transmodal association cortex retains selective structural influences. These findings provide new insight into how anatomical networks shape temporal processing in the human brain and suggest that intrinsic timescales emerge through distinct modes of interaction between structural constraints and functional dynamics.

## INTRODUCTION

Understanding how structural connectivity shapes functional brain dynamics remains a central challenge in network neuroscience (Deco et al., 2011; Fotiadis et al., 2024; Honey et al., 2009; Park & Friston, 2013; Schmitt, 2025; Sporns et al., 2002; Sporns et al., 2000). A widely used concept in this context is structure–function coupling (SFC), which quantifies the extent to which functional interactions between brain regions reflect the underlying anatomical connectivity derived from white-matter pathways (Bashirgonbadi et al., 2025; Baum et al., 2020; Popp et al., 2025; Preti & Van De Ville, 2019). Numerous neuroimaging studies have demonstrated that stronger structure–function coupling, measured at whole-brain or network scales, is associated with superior cognitive performance, including general intelligence and executive efficiency (Bashirgonbadi et al., 2025; Mišić et al., 2016; Popp et al., 2025; Preti & Van De Ville, 2019; Zimmermann et al., 2016). These findings suggest that the alignment between structure and function plays a fundamental role in effective brain computation.

Beyond static functional interactions, increasing attention has been devoted to the temporal organization of brain activity (Lurie et al., 2024; Munn et al., 2024; Northoff et al., 2025; van Es et al., 2025). In particular, intrinsic functional timescales have emerged as a key descriptor of how neural activity fluctuates over time. Functional timescales are commonly quantified using autocorrelation-based metrics or spectral properties of resting-state fMRI BOLD signals (Foster et al., 2016; Murray et al., 2014). Regions with short intrinsic timescales exhibit fast, transient dynamics dominated by higher-frequency fluctuations, whereas regions with long timescales display slow, temporally integrated activity dominated by low-frequency components.

Importantly, intrinsic timescales are not uniformly distributed across the cortex. Converging evidence from human fMRI (Mecklenbrauck et al., 2024; Raut et al., 2020), electrophysiology (Gao et al., 2020), and cross-species studies (Li et al., 2025; Song et al., 2024) has revealed a hierarchical organization, in which unimodal sensory and motor cortices exhibit short timescales, while transmodal association regions—including default mode and frontoparietal networks—show progressively longer timescales (Golesorkhi et al., 2021; Hasson et al., 2008; Huntenburg et al., 2018; Murray et al., 2014). This hierarchy closely mirrors functional specialization: unimodal regions support rapid stimulus-driven processing, whereas transmodal regions enable integrative and cognitively demanding operations that require information accumulation over extended temporal windows (Chaudhuri et al., 2015; Gao et al., 2020; Honey et al., 2012; Raut et al., 2020).

Several studies have linked this temporal hierarchy to regional structural connectivity strength (SC-strength). Empirical evidence indicates that brain regions with stronger or denser structural connections tend to exhibit slower functional dynamics and longer intrinsic timescales (Demirtaş et al., 2019; Liégeois et al., 2019). From a dynamical systems perspective, increased recurrent or long-range structural coupling can stabilize neural activity and promote temporal integration, leading to slower BOLD fluctuations. These findings suggest that SC-strength provides a mechanistic substrate through which anatomical connectivity influences regional functional timescales (Fallon et al., 2020; Gao et al., 2020; Ponce-Alvarez, 2025).

However, SC-strength alone does not fully explain the spatial organization of brain dynamics. The degree to which structural connectivity constrains functional interactions varies systematically across the cortex. Prior work has shown that structure–function coupling is strongest in unimodal sensory and motor regions, where functional connectivity closely follows anatomical pathways, and weakest in transmodal association cortex, where functional interactions increasingly diverge from structural constraints (Baum et al., 2020; Mišić et al., 2016; Preti & Van De Ville, 2019). This progressive decoupling has been interpreted as a marker of functional flexibility, enabling higher-order cognitive processes that are less rigidly bound to fixed anatomical circuits.

Notably, this spatial gradient of structure–function coupling parallels the hierarchy of intrinsic timescales. Unimodal regions combine high structure–function coupling with short timescales, reflecting fast, structure-driven processing (Vázquez-Rodríguez et al., 2019). In contrast, transmodal regions exhibit lower coupling and longer timescales, consistent with slow, integrative, and context-dependent dynamics (Gao et al., 2020). Together, these observations point to a close interaction between structural constraints, functional alignment, and temporal integration across the cortical hierarchy.

Integrating these findings suggests that structure–function coupling may mediate the relationship between SC-strength and functional timescales. In regions with strong coupling, anatomical connectivity may directly shape temporal dynamics. Conversely, in transmodal regions with weaker coupling, long intrinsic timescales may emerge from distributed and flexible functional interactions that are only partially constrained by structural architecture. This perspective also offers a refined interpretation of prior reports linking structure–function coupling to cognitive ability. While higher global coupling has been associated with improved cognitive performance (Popp et al., 2025; Zimmermann et al., 2016), regional specialization indicates that cognitive complexity relies on selective decoupling in the association cortex, accompanied by prolonged temporal integration.

From a signal processing perspective, functional brain activity can be modeled as a graph signal defined on the structural connectome (Huang et al., 2018; Medaglia et al., 2018; Preti & Van De Ville, 2019). Graph signal processing (GSP) provides a principled framework for decomposing functional signals into structure-aligned (coupled) and structure-liberal (decoupled) components using graph spectral modes (Bashirgonbadi et al., 2025; Ortega et al., 2018). Within this framework, structure–function coupling corresponds to signal smoothness with respect to the structural graph, while intrinsic timescales emerge from the temporal statistics of graph-filtered signals. This formulation provides a natural link between anatomical connectivity, temporal dynamics, and cognitive function.

Building on this framework, the present study investigates how structural connectivity, structure–function coupling, and intrinsic functional timescales jointly shape large-scale human brain dynamics. We test the hypothesis that intrinsic timescales derived from structure-coupled (graph low-pass) components of BOLD activity are more strongly associated with regional SC-strength than those derived from structure-decoupled (graph high-pass) components. We further hypothesize that the canonical structure–timescale relationship can be largely reproduced using structure-aligned signal components alone, whereas structure-decoupled activity exhibits weak dependence on anatomical connectivity. Extending these predictions, we examine whether association cortical regions display relatively longer intrinsic timescales within structure-decoupled signals compared to unimodal sensory regions. Finally, we assess the specificity of these effects using null models based on randomized graph spectral bases to determine whether observed relationships depend on the empirical organization of the structural connectome.

### GSP Methodology

Graph Signal Processing (GSP) provides a mathematical framework for analyzing signals defined on irregular domains, such as brain networks, by extending classical signal processing concepts to graph-structured data (Sandryhaila & Moura, 2013; Shuman et al., 2013; Zhao et al., 2023). Central to this framework are the Graph Fourier Transform (GFT) and graph-based filtering operations, which enable the decomposition of graph signals into spectral components. In the graph spectral domain, low-frequency components correspond to signals that vary smoothly over the underlying graph structure, whereas high-frequency components capture rapid spatial variations across nodes (Ortega et al., 2018; Yarandi & Babaie-Zadeh, 2023).

By applying the GFT, spatial patterns of brain activity can be examined at each time point relative to the topology of the structural connectome, providing a principled way to relate functional signals to anatomical connectivity (Cheng et al., 2023; Huang et al., 2018; Preti & Van De Ville, 2019). As such, GSP offers a powerful framework for investigating how functional brain dynamics unfold over structural networks and how network topology constrains spatial and temporal signal properties.

### Definition of Brain Graph and Graph Signal

The brain connectome is modeled as a weighted undirected graph *G* = (*v*, ***A***), where *v* = {1, 2, …, *N*} denotes the set of N nodes corresponding to cortical brain regions, and ***A*** ∈ ℝ^*N*×*N*^ is the weighted adjacency matrix. Each element ***A***_*i,j*_ quantifies the strength of the anatomical connection between regions i and j defined as the Number of Streamlines (NoS) obtained from white-matter tractography. These connectivity estimates are derived from diffusion-weighted imaging techniques such as Diffusion Spectrum Imaging (DSI) or Diffusion Tensor Imaging (DTI) (Catani et al., 2002).

From the adjacency matrix ***A***, the graph Laplacian is defined as ***L*** = ***D*** − ***A***, where ***D*** is the diagonal degree matrix with entries ***D***_*i,i*_ = ∑_*j*∈*v*_ ***A***_*i,j*_. The graph Laplacian encodes essential information about the topology of the structural network and serves as a fundamental operator for graph spectral analysis (Shuman et al., 2013).

Functional brain activity measured using functional Magnetic Resonance Imaging (fMRI) is represented as a graph signal **x** ∈ ℝ^N^ at each time point, where the i-th element corresponds to the BOLD signal of region i. Over time, functional activity is therefore represented by a matrix ***X*** ∈ ℝ^(N×T)^, with each column corresponding to a graph signal defined on the same structural graph.

Since the structural graph is undirected, the Laplacian matrix admits an eigendecomposition of the form (Leus et al., 2023; Mohammadi et al., 2023):

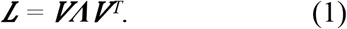

Here, ***Λ*** = diag (λ_0_, λ_1_, ⋯, λ_(N-1)_) contains the Laplacian eigenvalues ordered as λ_0_≤λ_1_≤⋯≤λ_(N-1)_, and ***V*** = [**v**_0_, **v**_1_,…, **v**_(N-1)_] denotes the corresponding orthonormal eigenvectors. The Graph Fourier Transform (GFT) of a graph signal **x** ∈ ℝ^N^ is defined as (Huang et al., 2018):

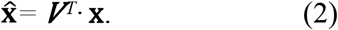

The original signal can be reconstructed via the inverse GFT as:

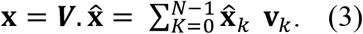

### Harmonic Frequency Extraction

The structural brain graph was constructed by parcellating the cortical gray matter into 360 regions using the Glasser atlas (Glasser et al., 2016), with each region representing a graph vertex. Edge weights were defined by the Number of Streamlines connecting each pair of regions, derived from diffusion tractography.

The graph Laplacian of this structural network was decomposed to obtain 360 eigenvectors, commonly referred to as graph harmonics, corresponding to the 360 cortical regions. Eigenvectors associated with smaller eigenvalues represent low-frequency graph modes that vary smoothly across the structural network, whereas eigenvectors associated with larger eigenvalues capture high-frequency spatial variations that are less aligned with the underlying anatomical connectivity (Huang et al., 2018; Ortega et al., 2018).

### Frequency Analysis of Functional Signals

For each time point, regional BOLD activity was averaged within each of the 360 regions of interest, yielding a graph signal defined on the structural connectome. The HCP resting-state fMRI dataset consists of 1200 time points, resulting in 1200 graph signals defined over an identical node set. To analyze functional dynamics in the graph spectral domain, each graph signal was projected onto the Laplacian eigenvectors using the GFT, yielding spectral coefficients that characterize the spatial frequency content of brain activity at each time point. To dissociate structure-aligned and structure-liberal components of functional activity, each graph signal was further decomposed into low-frequency (Coupled, **x**_*C*_) and high-frequency (Decoupled, **x**_*D*_) components using graph spectral filtering (Preti & Van De Ville, 2019). This decomposition enables the separation of functional dynamics that are smooth with respect to the structural network from those that exhibit high spatial variation across the connectome.

## Data and Analysis Methods

### Dataset

The Human Connectome Project (HCP) provides high-quality multimodal neuroimaging and behavioral data from a large cohort of healthy adults (Van Essen et al., 2013), offering a robust platform for investigating large-scale brain connectivity. In this study, we analyzed data from 100 unrelated healthy participants (54% female; mean age = 29.11 ± 3.67 years) with available resting-state fMRI (rs-fMRI) and diffusion-weighted imaging (DWI). Resting-state fMRI data were preprocessed using the HCP minimal preprocessing pipeline, which includes motion correction, distortion correction, intensity normalization, and projection to the 32k_fs_LR cortical surface space (Glasser et al., 2013). Structural connectivity matrices were derived from diffusion MRI using MRtrix3 (Tournier et al., 2019) employing anatomically constrained tractography, spherical deconvolution, and streamline filtering via SIFT2 to reduce tractography biases (Smith et al., 2015). Connectivity between 360 cortical regions, defined according to the Glasser atlas, was quantified using the number of streamlines. A group-level structural connectome was constructed using distance-dependent thresholding followed by logarithmic transformation to stabilize the distribution of edge weights. Analysis code is publicly available through the project GitHub repository: https://github.com/alibashir1364/Brain-Structure-Function-Coupling-and-BOLD-Dynamic.git.

Neuroimaging data were obtained from the https://db.humanconnectome.org. These resources are provided to facilitate transparency and reproducibility.

### Intrinsic BOLD Timescale Estimation

Intrinsic temporal properties of regional brain activity were quantified using the time-series analysis framework introduced by Fulcher et al. (Fallon et al., 2020). Timescales measures were computed separately for the original BOLD signal as well as for the structure-coupled and structure-decoupled components obtained via graph spectral filtering. For each regional time series, Relative Low-Frequency Power (RLFP) was calculated from the power spectral density as the proportion of spectral power below 0.14 Hz relative to the total spectral power, providing a robust index of slow spontaneous fluctuations (Sethi et al., 2017). Region-wise RLFP values were first computed at the subject level and subsequently averaged across participants. This procedure yielded intrinsic timescale estimates for the structure-coupled (RLFP_low_) and structure-decoupled (RLFP_high_) components of BOLD dynamics.

### Structure–Timescale Analysis and Statistical Testing

The association between intrinsic timescales and structural connectivity strength was assessed using partial Spearman correlation, controlling for regional volume to account for potential confounds related to parcel size. Correlation analyses were conducted at both the group level, using timescale measures averaged across participants, and at the inter-individual level by computing correlations separately for each participant. Differences between structure-coupled and structure-decoupled components were evaluated using paired non-parametric statistical tests across participants, and effect sizes were quantified using Cohen’s d. To assess the specificity of observed effects to the empirical structural connectome, null models were generated by randomizing the graph Fourier basis while preserving the Laplacian eigenvalue spectrum. Observed correlation values were then compared against the corresponding null distributions to evaluate statistical significance.

### Network-Level Analysis

To examine regional specificity, cortical areas were categorized into transmodal and unimodal systems based on established cortical hierarchies. All analyses were repeated separately within each system, enabling direct comparison of structure–timescale coupling across distinct levels of cortical organization.

## Results

Figure 1 summarizes the overall analysis pipeline. Structural connectivity matrices derived from diffusion MRI define the underlying brain graph, on which resting-state BOLD activity is modeled as a graph signal. Graph spectral decomposition is then used to separate functional activity into structure-coupled (low-pass) and structure-decoupled (high-pass) components. Intrinsic temporal properties are subsequently quantified using established time-series measures and related to node-wise structural connectivity strength.

**Figure 1.**
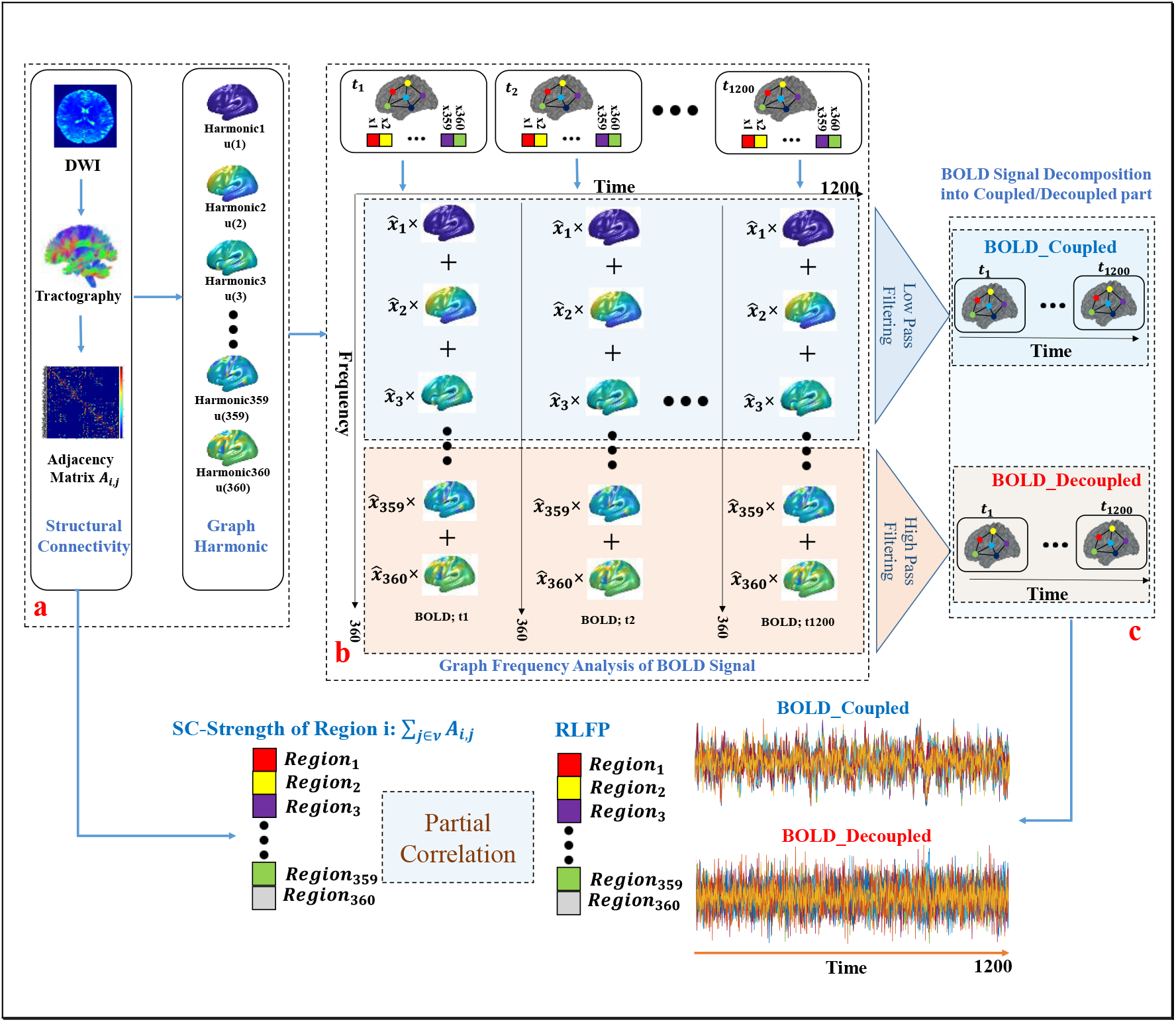
Workflow of the analytical pipeline. (Box a) details the construction of the structural connectivity matrix from DTI tractography, with structural graph eigenvectors (frequency harmonics) on the right. (Box b-top) shows the BOLD time series used as functional graph signals, with each time point represented as an individual graph signal (totaling 1200 signals, denoted t_1_, t_2_, …, t_1200_). (Box b-down) At each time point t_i_, the regional brain activity **x** is represented as a linear combination of graph harmonics through its spectral coefficients 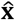. The spectral energy density of the activity is then computed, and a median-split criterion is applied to partition the graph spectrum into low- and high-frequency bands. This procedure enables the decomposition of brain activity into structurally **coupled** components (**x**_*c*_), capturing graph-smooth dynamics, and **decoupled** components (**x**_*d*_), reflecting graph-liberal variations. The vertical axis represents graph frequencies, indexed by the 360 eigenvectors (harmonics) of the graph Laplacian, ordered from low to high frequency. (Box c) The resulting coupled and decoupled BOLD signal components are defined on the empirical structural graph and span the full resting-state time series (t=1, …, 1200). Each component represents a time-resolved graph signal, preserving the original temporal resolution while selectively capturing dynamics that are either coupled with or decoupled from the underlying structural connectivity. For each cortical region, structural connectivity strength (SC strength) is computed from the empirical connectome. In parallel, intrinsic timescale measures (RLFP) are estimated for all regions separately from the coupled and decoupled BOLD components. To quantify the relationship between anatomical embedding and temporal dynamics, partial Spearman correlations are computed between SC strength and RLFP across regions while controlling for regional volume.

### Relationship between structural connectivity strength and intrinsic BOLD timescales

We first evaluated the baseline relationship between structural connectivity strength and intrinsic functional timescales using resting-state fMRI data from the Human Connectome Project. Structural connectivity strength was defined as the sum of streamline weights per region, while intrinsic timescales were quantified using the relative low-frequency power (RLFP) of regional BOLD signals. As shown in Figure 2-A, structural connectivity strength exhibited a clear positive association with RLFP across cortical regions, indicating that nodes with stronger structural embedding tend to exhibit slower temporal dynamics. To control for potential confounds related to parcel size, partial Spearman correlations were computed while regressing out regional volume. At the group level, this analysis revealed a significant positive association (r = 0.49, p = 9.49×e^−23^), consistent with prior findings in both human and animal studies (Fallon et al., 2020; Sethi et al., 2017). The spatial distribution of intrinsic timescales is illustrated in Figure 2-B, where RLFP values vary systematically across the cortex. Higher values are observed in association regions, while lower values are found in primary sensory and motor areas. This spatial heterogeneity reflects the well-established cortical hierarchy of temporal processing and provides a spatial context for the observed structure–timescale relationship (Foster et al., 2016; Golesorkhi et al., 2021; Lurie et al., 2024; Mecklenbrauck et al., 2024; Munn et al., 2024; Murray et al., 2014). Collectively, these results reproduce the canonical association between structural connectivity strength and intrinsic functional timescales, establishing a reference point for subsequent graph-spectral analyses.

**Figure 2.**
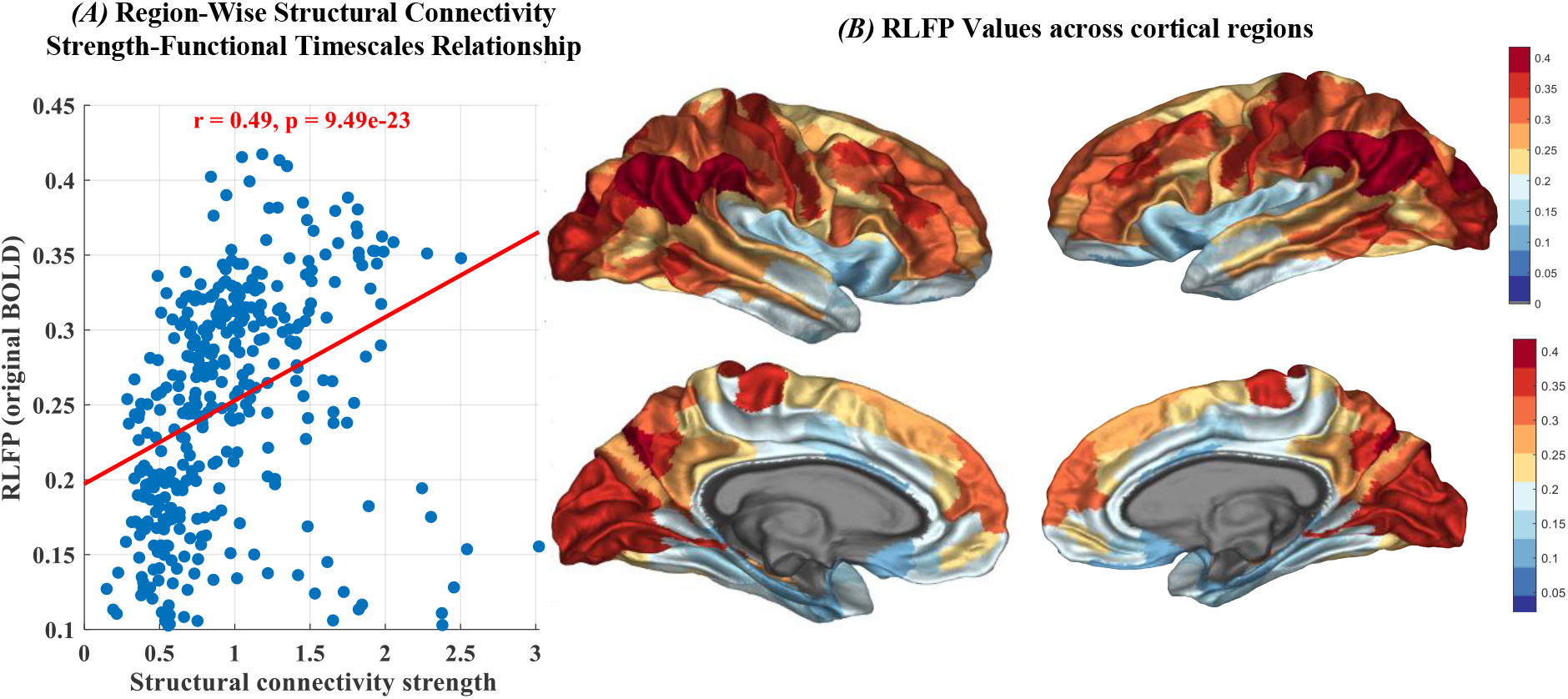
Structure–timescale relationship in the human cortex. (A) Scatter plot illustrating the relationship between structural connectivity strength and intrinsic functional timescales, quantified using relative low-frequency power (RLFP), across 360 cortical regions. Each point represents a brain region. (B) Spatial cortical map of RLFP values, revealing a heterogeneous distribution of intrinsic timescales across the cortex. Together, these results replicate prior findings linking structural connectivity strength to intrinsic functional dynamics using high-quality Human Connectome Project data.

### Structural connectivity selectively constrains graph-smooth functional dynamics

To determine whether the observed structure–timescale relationship depends on the alignment of functional activity with the structural graph, BOLD signals were decomposed into graph-coupled (low-pass) and graph-decoupled (high-pass) components using graph spectral filtering. Intrinsic timescales were then estimated separately for each component using RLFP. As shown in Figure 3-A, intrinsic timescales derived from the coupled (low-pass) component exhibited a strong positive association with structural connectivity strength. In contrast, the corresponding relationship for the decoupled (high-pass) component was substantially attenuated (Figure 3-B), indicating reduced sensitivity of structurally misaligned activity to the underlying graph topology. This dissociation was confirmed by group-level partial Spearman correlation analysis controlling for regional volume, which revealed a significant difference between the two components (Fisher Z = 4.86, p = 1.16e^−06^; Figure 3-C). Specifically, the correlation between structural connectivity strength and RLFP was significantly higher for the graph-coupled component than for the graph-decoupled component. These findings support Hypothesis H1, demonstrating that structural connectivity selectively constrains graph-smooth functional dynamics, and Hypothesis H3, indicating that this constraint is markedly reduced for graph-decoupled activity. Overall, these results show that the classical structure–timescale relationship is not a uniform property of functional signals, but is preferentially expressed in graph-smooth components aligned with the structural connectome.

**Figure 3.**
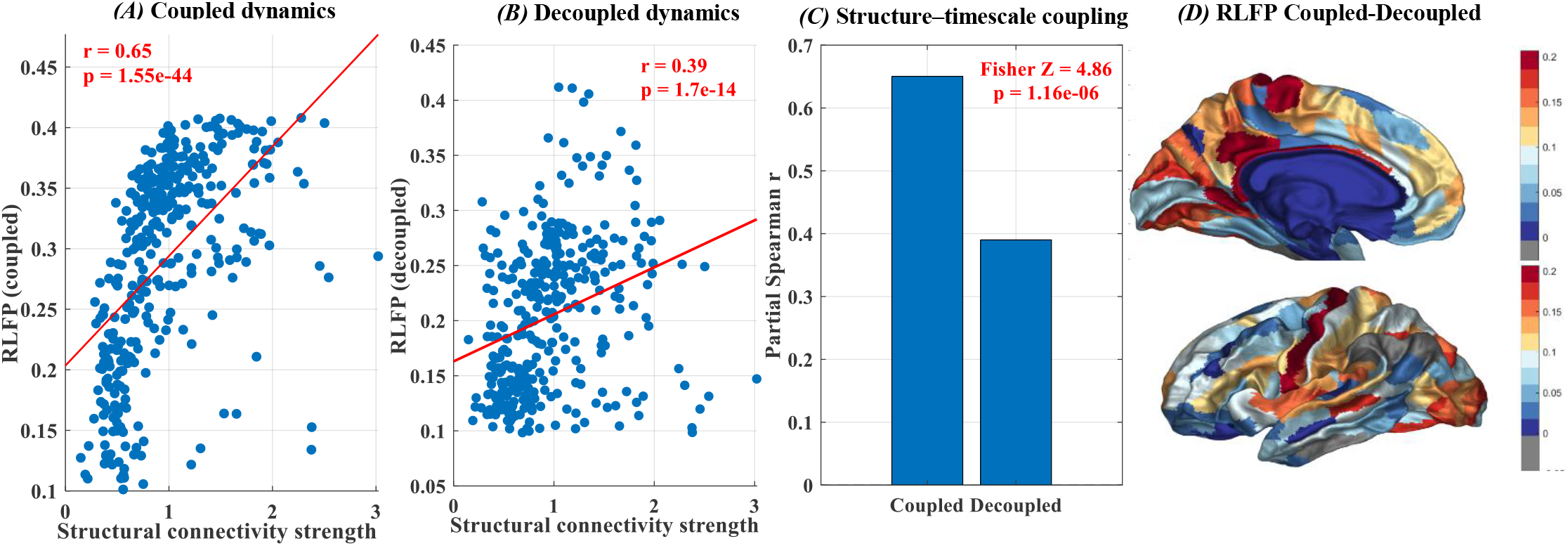
Structural connectivity selectively constrains graph-smooth BOLD signal. (A) Scatter plot showing the relationship between structural connectivity strength and intrinsic timescales estimated from the graph-coupled (low-pass) BOLD component across cortical regions. (B) Corresponding scatter plot for the graph-decoupled (high-pass) BOLD component. Group-level partial Spearman correlation coefficients quantifying the structure–timescale relationship for coupled and decoupled signals, controlling for regional volume. (D) Spatial cortical map illustrating the regional difference in intrinsic timescales between coupled and decoupled components (low-pass minus high-pass). These results demonstrate that structural connectivity strength preferentially constrains graph-smooth functional activity, while the relationship is markedly reduced for graph-decoupled dynamics.

### Inter-Individual Robustness of Structure–Timescale Coupling

To assess whether the observed dissociation reflects a robust individual-level phenomenon rather than a consequence of group averaging, structure–timescale coupling was evaluated independently for each subject. For each individual, partial Spearman correlations between structural connectivity strength and intrinsic timescales were computed separately for graph-coupled (low-pass) and graph-decoupled (high-pass) signals. As illustrated in Figure 4-A, subject-wise correlations consistently showed stronger positive associations for the coupled component relative to the decoupled component. While the magnitude of correlations varied across individuals, the direction of the effect was highly consistent. This consistency is further highlighted in Figure 4-B, where paired comparisons demonstrate that the majority of subjects exhibit higher correlations for coupled signals than for decoupled signals. A paired statistical test confirmed that this within-subject difference was highly significant (Wilcoxon signed-rank test, p = 3.9e^−18^). The corresponding effect size (Figure 4-C) indicates a medium-to-large effect, demonstrating that the observed dissociation is both statistically robust and practically meaningful. These findings support Hypothesis H4 and indicate that the preferential coupling between structural connectivity and intrinsic timescales is a stable and reproducible property across individuals, rather than an artifact of group-level averaging.

**Figure 4.**
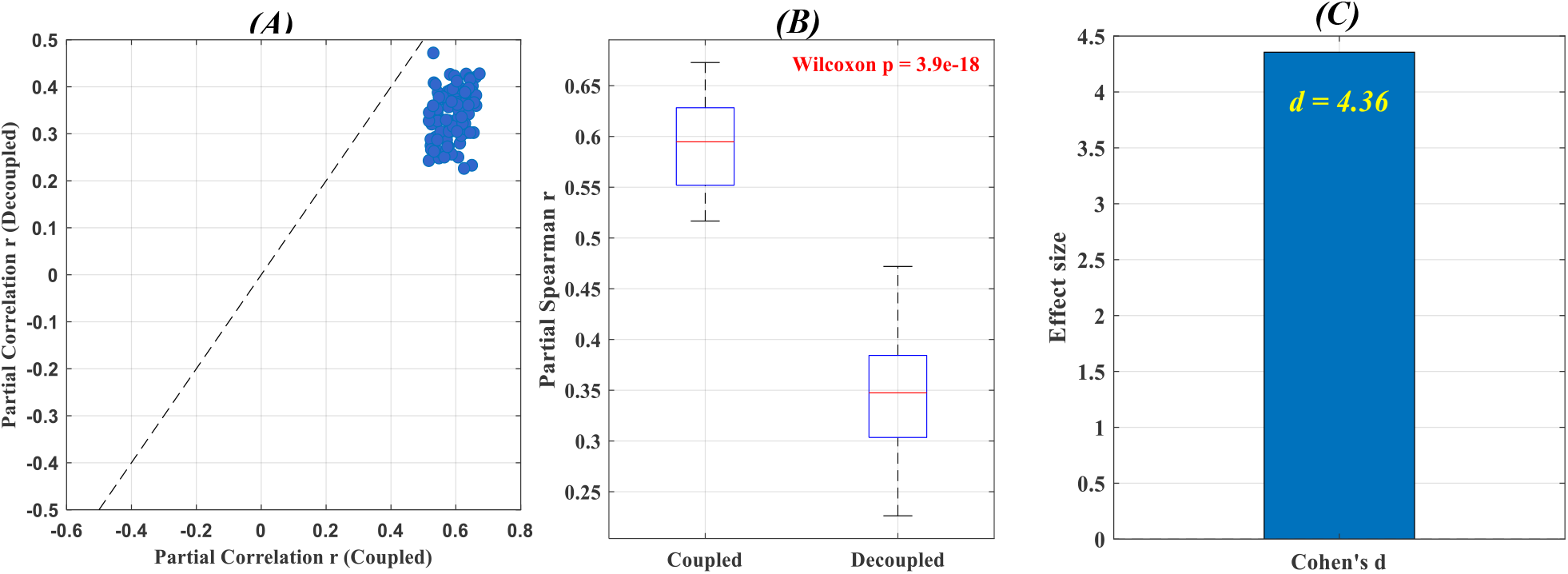
Inter-individual robustness of structure–timescale relationship. (A) Subject-wise partial Spearman correlation coefficients between structural connectivity strength and intrinsic timescales computed from graph-coupled (low-pass) and graph-decoupled (high-pass) BOLD components. Each point represents one subject. (B) Paired comparison of subject-level correlation values for coupled and decoupled components, illustrating a consistent shift toward stronger coupling in the graph-coupled signal. (C) Effect size (Cohen’s *d*) quantifying the magnitude of the within-subject difference between coupled and decoupled correlations. These results demonstrate that the preferential association between structural connectivity and intrinsic timescales is consistently observed across individuals and is not driven by group-level averaging.

### Null Model Validation of SC-Strength–Timescale Relationship

To evaluate whether the observed associations reflect intrinsic properties of the empirical structural connectome rather than generic spectral effects, we employed a null model based on randomized graph Fourier bases (Huang et al., 2018). Specifically, the Laplacian eigenvectors were randomized while preserving the original BOLD time series, and partial Spearman correlations between structural connectivity strength and intrinsic timescales were recomputed across null realizations. For the graph-coupled (low-pass) component, the observed correlation (r = 0.6503) lay far outside the null distribution (null mean = −0.0001 ± 0.0566), corresponding to a Z-score of 11.50 (p ≈ 0; Figure 5-A). This indicates that the strong association observed in the empirical data cannot be attributed to random spectral structure. A similar pattern was observed for the graph-decoupled (high-pass) component. The observed correlation (r = 0.3901) significantly exceeded the null distribution (null mean = −0.0003 ± 0.0530; Z = 7.37, p = 1.76 × 10^−13^; Figure 5-B). Although weaker than the coupled case, the relationship remained highly significant relative to the null model. Importantly, this analysis does not directly compare coupled and decoupled components, but instead establishes that, for each signal separately, the empirical structural graph imposes a statistically significant constraint on intrinsic timescales. These results confirm that the observed structure–timescale relationships depend critically on the true organization of the structural connectome and are not reducible to trivial spectral artifacts.

**Figure 5.**
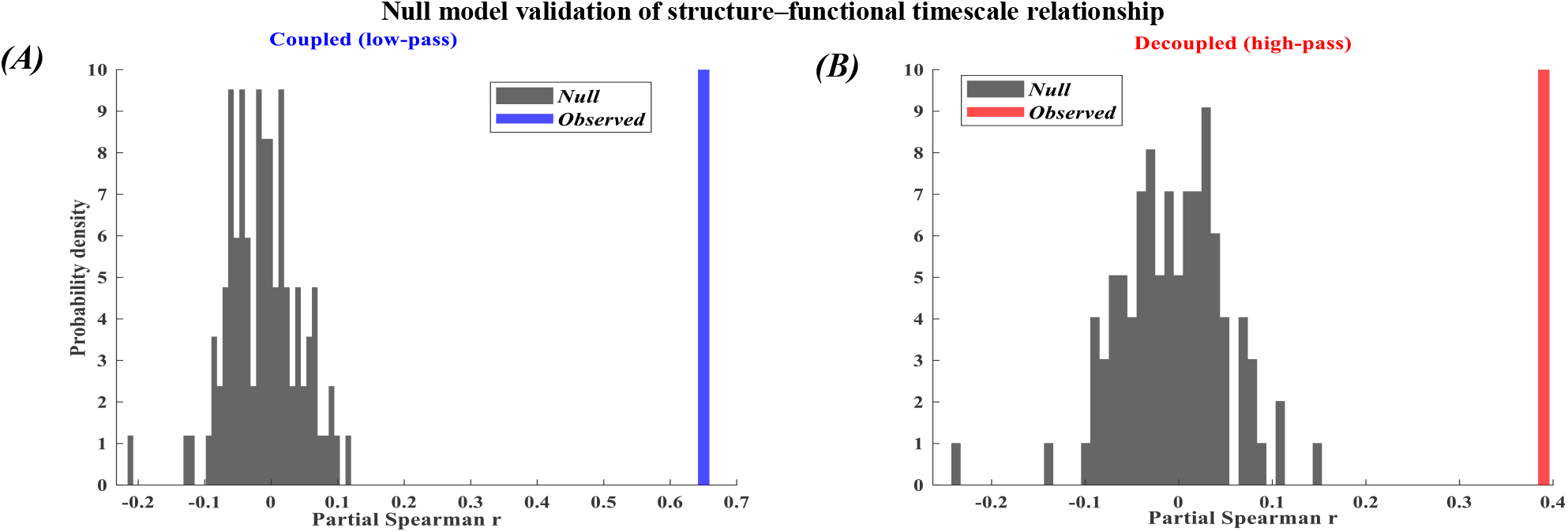
Null model validation of structure–functional timescale relationship. Null distributions of partial Spearman correlations between regional structural connectivity strength and intrinsic timescales were generated by randomizing structural connectivity weights while preserving matrix symmetry and controlling for regional volume. (A) Null distribution for the coupled (low-pass) signal, with the observed correlation obtained from the empirical connectome indicated by the vertical line. Corresponding null distribution for the decoupled (high-pass) signal. In both cases, randomization of structural connectivity markedly attenuates the structure–timescale association, and the observed correlations significantly deviate from the null distributions, demonstrating that the reported effects critically depend on the empirical structural connectome.

### Network-Specific Constraint of Decoupled Dynamics

To investigate whether structural constraints on decoupled functional dynamics vary across cortical hierarchies, analyses were performed separately for transmodal and unimodal regions. As shown in Figure 6-A, transmodal regions exhibited a stronger partial Spearman correlation between structural connectivity strength and intrinsic timescales of graph-decoupled (high-pass) signals compared to unimodal regions, after controlling for regional volume. Group-level comparisons (Figure 6-B) further demonstrate differences in the mean RLFP of decoupled dynamics across cortical systems. A Fisher Z-test confirmed that the strength of the structure–timescale relationship differed significantly between transmodal and unimodal cortex (Fisher Z = 2.33; p = 0.0196), indicating a hierarchy-dependent modulation of structural constraints. Regional scatter plots (Figure 6-C and Figure 6-D) illustrate these effects more directly. Transmodal regions show a clearer positive relationship between structural connectivity strength and decoupled intrinsic timescales, whereas unimodal regions exhibit greater dispersion and a weaker linear trend. These findings indicate that even graph-decoupled dynamics are not entirely unconstrained, but remain selectively influenced by structural connectivity in higher-order association cortex.

**Figure 6.**
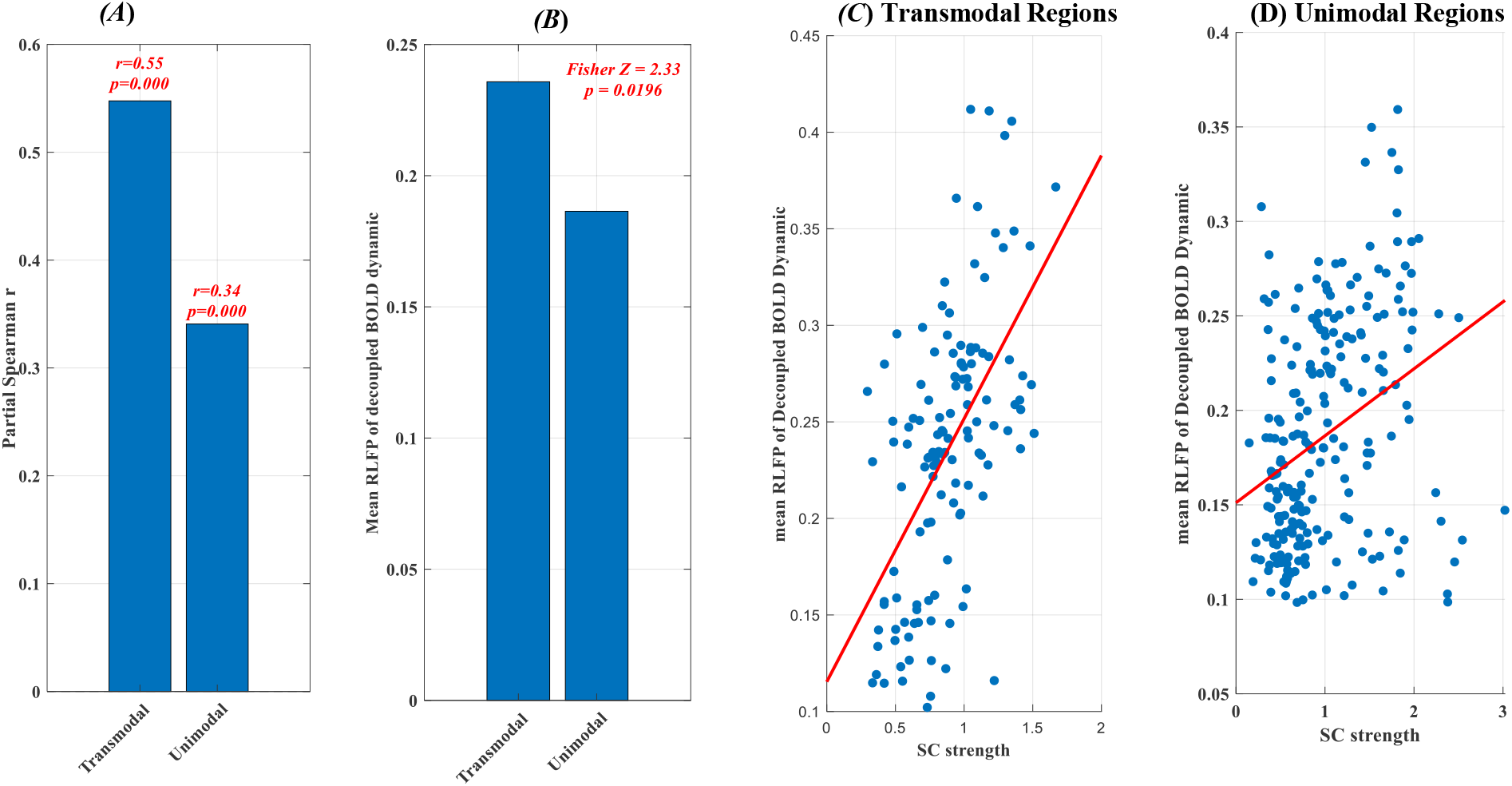
Hierarchy-dependent structural constraint of decoupled functional dynamics. (A) Partial Spearman correlations between SC-strength and intrinsic timescales of graph-decoupled (high-pass) BOLD signals (high-pass), shown separately for transmodal and unimodal cortex while controlling for regional volume. (B) Mean RLFP of graph-decoupled functional dynamics across cortical groups, with the between-group difference in correlation strength assessed using a Fisher Z-test. (C) Scatter plot illustrating the regional relationship between SC-strength and RLFP of graph-decoupled functional dynamics in transmodal cortex. (D) Corresponding scatter plot for unimodal cortex. Together, these results demonstrate that structural connectivity more strongly constrains decoupled functional dynamics in transmodal cortex than in unimodal regions.

## Discussion

In this study, we leveraged a graph signal processing (GSP) framework to examine how the structural connectome constrains the temporal statistics of functional signals defined on it. By modeling resting-state BOLD activity as graph signals and decomposing them into graph-spectral components, we show that the canonical association between structural connectivity strength (SC-strength) and intrinsic functional timescales is primarily carried by graph-smooth (low-frequency) activity aligned with the structural network. This result provides a signal-processing interpretation of prior observations linking anatomical connectivity to slow cortical dynamics, indicating that structure–timescale coupling is fundamentally a graph-spectral phenomenon.

Within the GSP formulation, low-frequency graph components correspond to signals that are smooth with respect to the structural Laplacian, reflecting coordinated activity across strongly connected regions. The strong association observed between these components and intrinsic timescales suggests that temporally slow dynamics preferentially emerge in signal subspaces aligned with the structural graph. This is consistent with theoretical models in which dense or recurrent connectivity promotes temporal integration and stabilizes network dynamics, leading to slower fluctuations (Honey et al., 2012). In contrast, high-frequency graph components represent signals with rapid spatial variation across the network. The substantially weaker relationship between these components and SC-strength indicates that structurally misaligned activity is less directly constrained by the underlying graph topology.

Importantly, the separation between structure-coupled and structure-decoupled dynamics is not limited to group-level analyses, but is consistently observed at the level of individual subjects. This inter-individual consistency indicates that the preferential coupling between structural connectivity strength and graph-smooth temporal dynamics reflects a stable property of networked brain signals, rather than an artifact of aggregation. This inter-individual robustness strengthens the interpretation that structural connectivity exerts a selective and reproducible influence on intrinsic neural timescales through graph-smooth activity.

Our results refine the notion of structure–function decoupling in higher-order cortex. While structure-decoupled signals are often interpreted as reflecting dynamics that are released from anatomical constraints, our findings suggest a more nuanced picture. Specifically, transmodal regions exhibited significantly elevated intrinsic timescales in the decoupled component, alongside a structure–timescale relationship that differed markedly from that of the unimodal cortex. Importantly, this relationship was not abolished, but instead manifested as a distinct mode of structural modulation. This observation indicates that the transmodal cortex does not operate in a structure-independent regime, but rather engages structural connectivity in a qualitatively different manner. In these regions, anatomical connectivity may act as an enabling scaffold that supports the emergence of slow, integrative dynamics, rather than directly constraining local temporal properties. Such a facilitative role of structure is consistent with the integrative function of transmodal cortex and aligns with theoretical accounts in which higher-order regions exploit distributed connectivity to sustain extended temporal processing windows. In contrast, unimodal regions exhibited weaker and less differentiated structure–timescale relationships in the decoupled signal, consistent with more locally driven dynamics and reduced reliance on large-scale integration. Together, these findings support a hierarchical view of structure–function interaction in which the impact of anatomical connectivity on intrinsic dynamics varies across cortical systems, rather than uniformly diminishing along the cortical hierarchy.

The specificity of our findings to the empirical structural connectome was confirmed using graph-based null models. Randomization of the graph Fourier basis, while preserving the spectral properties of the Laplacian, substantially reduced the observed structure–timescale relationships. This result demonstrates that the effects reported here are not a trivial consequence of spectral filtering or low-frequency dominance in BOLD signals, but instead depend critically on the spatial organization of white-matter connectivity.

Together, these findings advance our understanding of structure–function relationships by demonstrating that structural connectivity does not uniformly constrain all aspects of spontaneous neural activity. Rather, it selectively shapes the temporal properties of graph-smooth activity, while allowing for regionally specific structure-decoupled dynamics, particularly in association cortex. This spectral dissociation provides a principled framework for reconciling anatomical constraints with functional flexibility in large-scale brain dynamics.

By integrating graph signal processing with intrinsic timescale analysis, this study provides a mechanistic account of how the brain’s structural scaffold shapes spontaneous neural dynamics. Structural connectivity selectively constrains graph-smooth activity to produce hierarchical intrinsic timescales, while allowing structure-decoupled fluctuations that are particularly prominent in association cortex. These findings highlight the utility of graph spectral approaches for dissecting the multiscale organization of brain dynamics and offer a principled framework for future investigations of structure–function relationships.

## Acknowledgment

In the preparation of this article, artificial intelligence (AI) tools were utilized to assist in various stages of the writing process. All AI-generated content was rigorously reviewed, verified, and edited by the authors to ensure accuracy, originality, and alignment with academic standards. Responsibility for the final content rests solely with the authors.

## Notes

https://github.com/alibashir1364/Brain-Structure-Function-Coupling-and-BOLD-Dynamic.git

### Competing Interest Statement

The authors have declared no competing interest.

### Summary of Updates

This revision corrects the author order due to an error in the author metadata of the previous version. The manuscript content, including the analyses, results, and conclusions, remains unchanged.

## References

Bashirgonbadi, A., Salehi, M. R., & Soltanian-Zadeh, H. (2025). Vertex-Frequency Analysis of Brain Networks: Unveiling the Connection Between Structure-Function Coupling and Cognitive Ability. IEEE Transactions on Signal and Information Processing over Networks, 11, 1582–1591.

Baum, G. L., Cui, Z., Roalf, D. R., Ciric, R., Betzel, R. F., Larsen, B., Cieslak, M., Cook, P. A., Xia, C. H., & Moore, T. M. (2020). Development of structure–function coupling in human brain networks during youth. Proceedings of the National Academy of Sciences, 117(1), 771–778.

Catani, M., Howard, R. J., Pajevic, S., & Jones, D. K. (2002). Virtual in vivo interactive dissection of white matter fasciculi in the human brain. NeuroImage, 17(1), 77–94.

Chaudhuri, R., Knoblauch, K., Gariel, M.-A., Kennedy, H., & Wang, X.-J. (2015). A large-scale circuit mechanism for hierarchical dynamical processing in the primate cortex. Neuron, 88(2), 419–431.

Cheng, C., Chen, Y., Lee, Y. J., & Sun, Q. (2023). SVD-based graph Fourier transforms on directed product graphs. IEEE Transactions on Signal and Information Processing over Networks.

Deco, G., Jirsa, V. K., & McIntosh, A. R. (2011). Emerging concepts for the dynamical organization of resting-state activity in the brain. Nature reviews neuroscience, 12(1), 43–56.

Demirtaş, M., Burt, J. B., Helmer, M., Ji, J. L., Adkinson, B. D., Glasser, M. F., Van Essen, D. C., Sotiropoulos, S. N., Anticevic, A., & Murray, J. D. (2019). Hierarchical heterogeneity across human cortex shapes large-scale neural dynamics. Neuron, 101(6), 1181–1194. e1113.

Fallon, J., Ward, P. G., Parkes, L., Oldham, S., Arnatkevičiūtė, A., Fornito, A., & Fulcher, B. D. (2020). Timescales of spontaneous fMRI fluctuations relate to structural connectivity in the brain. Network Neuroscience, 4(3), 788–806.

Foster, B. L., He, B. J., Honey, C. J., Jerbi, K., Maier, A., & Saalmann, Y. B. (2016). Spontaneous neural dynamics and multi-scale network organization. Frontiers in systems neuroscience, 10, 7.

Fotiadis, P., Parkes, L., Davis, K. A., Satterthwaite, T. D., Shinohara, R. T., & Bassett, D. S. (2024). Structure–function coupling in macroscale human brain networks. Nature reviews neuroscience, 25(10), 688–704.

Gao, R., Van den Brink, R. L., Pfeffer, T., & Voytek, B. (2020). Neuronal timescales are functionally dynamic and shaped by cortical microarchitecture. elife, 9, e61277.

Glasser, M. F., Coalson, T. S., Robinson, E. C., Hacker, C. D., Harwell, J., Yacoub, E., Ugurbil, K., Andersson, J., Beckmann, C. F., & Jenkinson, M. (2016). A multi-modal parcellation of human cerebral cortex. Nature, 536(7615), 171–178.

Glasser, M. F., Sotiropoulos, S. N., Wilson, J. A., Coalson, T. S., Fischl, B., Andersson, J. L., Xu, J., Jbabdi, S., Webster, M., & Polimeni, J. R. (2013). The minimal preprocessing pipelines for the Human Connectome Project. NeuroImage, 80, 105–124.

Golesorkhi, M., Gomez-Pilar, J., Tumati, S., Fraser, M., & Northoff, G. (2021). Temporal hierarchy of intrinsic neural timescales converges with spatial core-periphery organization. Communications biology, 4(1), 277.

Hasson, U., Yang, E., Vallines, I., Heeger, D. J., & Rubin, N. (2008). A hierarchy of temporal receptive windows in human cortex. Journal of Neuroscience, 28(10), 2539–2550.

Honey, C. J., Sporns, O., Cammoun, L., Gigandet, X., Thiran, J.-P., Meuli, R., & Hagmann, P. (2009). Predicting human resting-state functional connectivity from structural connectivity. Proceedings of the National Academy of Sciences, 106(6), 2035–2040.

Honey, C. J., Thesen, T., Donner, T. H., Silbert, L. J., Carlson, C. E., Devinsky, O., Doyle, W. K., Rubin, N., Heeger, D. J., & Hasson, U. (2012). Slow cortical dynamics and the accumulation of information over long timescales. Neuron, 76(2), 423–434.

Huang, W., Bolton, T. A. W., Medaglia, J. D., Bassett, D. S., Ribeiro, A., & Van De Ville, D. (2018). A Graph Signal Processing Perspective on Functional Brain Imaging. Proceedings of the IEEE, 106(5), 868–885. 10.1109/JPROC.2018.2798928

Huntenburg, J. M., Bazin, P.-L., & Margulies, D. S. (2018). Large-scale gradients in human cortical organization. Trends in Cognitive Sciences, 22(1), 21–31.

Leus, G., Marques, A. G., Moura, J. M., Ortega, A., & Shuman, D. I. (2023). Graph Signal Processing: History, development, impact, and outlook. IEEE signal processing magazine, 40(4), 49–60.

Li, G., Li, S., & Wang, X.-J. (2025). A hierarchy of time constants and reliable signal propagation in the marmoset cerebral cortex. Nature communications.

Liégeois, R., Li, J., Kong, R., Orban, C., Van De Ville, D., Ge, T., Sabuncu, M. R., & Yeo, B. T. (2019). Resting brain dynamics at different timescales capture distinct aspects of human behavior. Nature communications, 10(1), 2317.

Lurie, D. J., Pappas, I., & D’Esposito, M. (2024). Cortical timescales and the modular organization of structural and functional brain networks. Human Brain Mapping, 45(2), e26587.

Mecklenbrauck, F., Sepulcre, J., Fehring, J., & Schubotz, R. I. (2024). Decoding cortical chronotopy—Comparing the influence of different cortical organizational schemes. NeuroImage, 303, 120914.

Medaglia, J. D., Huang, W., Karuza, E. A., Kelkar, A., Thompson-Schill, S. L., Ribeiro, A., & Bassett, D. S. (2018). Functional alignment with anatomical networks is associated with cognitive flexibility. Nature human behaviour, 2(2), 156–164.

Mišić, B., Betzel, R. F., De Reus, M. A., Van Den Heuvel, M. P., Berman, M. G., McIntosh, A. R., & Sporns, O. (2016). Network-level structure-function relationships in human neocortex. Cerebral Cortex, 26(7), 3285–3296.

Mohammadi, S., Babaie-Zadeh, M., & Thanou, D. (2023). Graph signal separation based on smoothness or sparsity in the frequency domain. IEEE Transactions on Signal and Information Processing over Networks, 9, 152–161.

Munn, B. R., Müller, E. J., Favre-Bulle, I., Scott, E., Lizier, J. T., Breakspear, M., & Shine, J. M. (2024). Multiscale organization of neuronal activity unifies scale-dependent theories of brain function. Cell, 187(25), 7303–7313. e7315.

Murray, J. D., Bernacchia, A., Freedman, D. J., Romo, R., Wallis, J. D., Cai, X., Padoa-Schioppa, C., Pasternak, T., Seo, H., & Lee, D. (2014). A hierarchy of intrinsic timescales across primate cortex. Nature neuroscience, 17(12), 1661–1663.

Northoff, G., Wolman, A., & Zhang, J. (2025). Brain dynamics shape cognition–Spatiotemporal Neuroscience. Physics of Life Reviews.

Ortega, A., Frossard, P., Kovačević, J., Moura, J. M., & Vandergheynst, P. (2018). Graph signal processing: Overview, challenges, and applications. Proceedings of the IEEE, 106(5), 808–828.

Park, H.-J., & Friston, K. (2013). Structural and functional brain networks: from connections to cognition. Science, 342(6158), 1238411.

Ponce-Alvarez, A. (2025). Network Mechanisms Underlying the Regional Diversity of Variance and Time Scales of the Brain’s Spontaneous Activity Fluctuations. Journal of Neuroscience, 45(10).

Popp, J. L., Thiele, J. A., Faskowitz, J., Seguin, C., Sporns, O., & Hilger, K. (2025). Structural-functional brain network coupling during cognitive demand reveals intelligence-relevant communication strategies. Communications biology, 8(1), 855.

Preti, M. G., & Van De Ville, D. (2019). Decoupling of brain function from structure reveals regional behavioral specialization in humans. Nature communications, 10(1), 4747.

Raut, R. V., Snyder, A. Z., & Raichle, M. E. (2020). Hierarchical dynamics as a macroscopic organizing principle of the human brain. Proceedings of the National Academy of Sciences, 117(34), 20890–20897.

Sandryhaila, A., & Moura, J. M. (2013). Discrete signal processing on graphs. IEEE transactions on signal processing, 61(7), 1644–1656.

Schmitt, O. (2025). Relationships and representations of brain structures, connectivity, dynamics and functions. Progress in Neuro-Psychopharmacology and Biological Psychiatry, 138, 111332.

Sethi, S. S., Zerbi, V., Wenderoth, N., Fornito, A., & Fulcher, B. D. (2017). Structural connectome topology relates to regional BOLD signal dynamics in the mouse brain. Chaos: An interdisciplinary journal of nonlinear science, 27(4).

Shuman, D. I., Narang, S. K., Frossard, P., Ortega, A., & Vandergheynst, P. (2013). The emerging field of signal processing on graphs: Extending high-dimensional data analysis to networks and other irregular domains. IEEE signal processing magazine, 30(3), 83–98.

Smith, R. E., Tournier, J.-D., Calamante, F., & Connelly, A. (2015). SIFT2: Enabling dense quantitative assessment of brain white matter connectivity using streamlines tractography. NeuroImage, 119, 338–351.

Song, M., Shin, E. J., Seo, H., Soltani, A., Steinmetz, N. A., Lee, D., Jung, M. W., & Paik, S.-B. (2024). Hierarchical gradients of multiple timescales in the mammalian forebrain. Proceedings of the National Academy of Sciences, 121(51), e2415695121.

Sporns, O., Tononi, G., & Edelman, G. (2002). Theoretical neuroanatomy and the connectivity of the cerebral cortex. Behavioural brain research, 135(1-2), 69–74.

Sporns, O., Tononi, G., & Edelman, G. M. (2000). Theoretical neuroanatomy: relating anatomical and functional connectivity in graphs and cortical connection matrices. Cerebral Cortex, 10(2), 127–141.

Tournier, J.-D., Smith, R., Raffelt, D., Tabbara, R., Dhollander, T., Pietsch, M., Christiaens, D., Jeurissen, B., Yeh, C.-H., & Connelly, A. (2019). MRtrix3: A fast, flexible and open software framework for medical image processing and visualisation. NeuroImage, 202, 116137.

van Es, M. W., Higgins, C., Gohil, C., Quinn, A. J., Vidaurre, D., & Woolrich, M. W. (2025). Large-scale cortical functional networks are organized in structured cycles. Nature neuroscience, 28(10), 2118–2128.

Van Essen, D. C., Smith, S. M., Barch, D. M., Behrens, T. E., Yacoub, E., Ugurbil, K., & Consortium, W.-M. H. (2013). The WU-Minn human connectome project: an overview. NeuroImage, 80, 62–79.

Vázquez-Rodríguez, B., Suárez, L. E., Markello, R. D., Shafiei, G., Paquola, C., Hagmann, P., Van Den Heuvel, M. P., Bernhardt, B. C., Spreng, R. N., & Misic, B. (2019). Gradients of structure–function tethering across neocortex. Proceedings of the National Academy of Sciences, 116(42), 21219–21227.

Yarandi, M.-H. A., & Babaie-Zadeh, M. (2023). A closed-form solution for graph signal separation based on smoothness. IEEE Transactions on Signal and Information Processing over Networks, 9, 823–824.

Zhao, H., Xiang, W., & Lv, S. (2023). A variable parameter LMS algorithm based on generalized maximum correntropy criterion for graph signal processing. IEEE Transactions on Signal and Information Processing over Networks, 9, 140–151.

Zimmermann, J., Ritter, P., Shen, K., Rothmeier, S., Schirner, M., & McIntosh, A. R. (2016). Structural architecture supports functional organization in the human aging brain at a regionwise and network level. Human Brain Mapping, 37(7), 2645–2661.

